# A Parameter-free Deep Embedded Clustering Method for Single-cell RNA-seq Data

**DOI:** 10.1101/2021.12.19.473334

**Authors:** Yuansong Zeng, Zhuoyi Wei, Fengqi Zhong, Zixiang Pan, Yutong Lu, Yuedong Yang

**Affiliations:** School of Computer Science and Engineering, Sun Yat-sen University, Guangzhou 510000, China; Key Laboratory of Machine Intelligence and Advanced Computing (MOE), Guangzhou 510000, China

**Keywords:** single cell RNA sequencing, single cell clustering, Estimating the number of cell clusters, Dip-test, Deep embedded clustering

## Abstract

Clustering analysis is widely utilized in single-cell RNA-sequencing (scRNA-seq) data to discover cell heterogeneity and cell states. While many clustering methods have been developed for scRNA-seq analysis, most of these methods require to provide the number of clusters. However, it is not easy to know the exact number of cell types in advance, and experienced determination is not always reliable. Here, we have developed ADClust, an **a**utomatic **d**eep embedding **clust**ering method for scRNA-seq data, which can accurately cluster cells without requiring a predefined number of clusters. Specifically, ADClust first obtains low-dimensional representation through pre-trained autoencoder, and uses the representations to cluster cells into initial micro-clusters. The clusters are then compared in between by a statistical test, and similar micro-clusters are merged into larger clusters. According to the clustering, cell representations are updated so that each cell will be pulled toward centres of its assigned cluster and similar clusters, while cells are separated to keep distances between clusters. This is accomplished through jointly optimizing the carefully designed clustering and autoencoder loss functions. This merging process continues until convergence. ADClust was tested on eleven real scRNA-seq datasets, and shown to outperform existing methods in terms of both clustering performance and the accuracy on the number of the determined clusters. More importantly, our model provides high speed and scalability for large datasets.

## 1. Introduction

Recent advances in single cell RNA sequencing (scRNA-seq) technologies have paved the way for researchers to generate high-throughput single-cell gene expression [1]. A full characterization of transcriptome profiling at single-cell resolution holds enormous potential for discovering trajectories of different cell developmental states and investing the cellular heterogeneity [2, 3]. One important step to discover cell heterogeneity and cell states is to perform clustering analysis, which aims to group a set of cells into meaningful cell populations based on their transcriptome similarity [4, 5]. The clustering can be used as additional downstream analysis and provide a reference to build a cell atlas [6-8]. Nevertheless, the clustering is meeting grand challenges due to the characteristic of scRNA-seq data, such as sparsity and high dimensional features [9, 10].

To resolve these challenges, a wide variety of clustering algorithms have been developed for scRNA-seq analysis [4, 5]. Early popular algorithms are variants of K-means that divide cells into K clusters with K as the pre-determined cluster number. For example, scDeepCluster [11] uses K-means to obtain initial centres of clusters, and then pushes each cell to its most similar centres iteratively. Similar strategies have also been used by other methods like SAIC [12], scVDMC [13], and DESC [14]. On the other hand, graph clustering is based on the community detection algorithms that cluster neighbored cells based on a resolution parameter. For example, Seurat [15], one of the most widely used toolkits for scRNA-seq analysis, connects cells into a KNN-graph and then partitions the graph into communities (clusters) through a predetermined resolution parameter, where a higher resolution generates a greater number of clusters. Similar strategies have also been used by other methods like SNN-Cliq [16] and SCANPY[17]. While these two classes of methods are robust, they need a parameter (K or resolution) as a priori, which unfortunately is seldom known in advance.

To avoid the predetermined parameter, SIMLR[18] pre-estimates the cluster number as the rank constraint and then combines graph diffusion to learn a cell similarity measure for clustering. Similarly, SC3[19] also pre-estimates the cluster number, but it then uses K-means to cluster cells from different eigenvectors, and constructs a consensus matrix for clustering. However, the pre-estimated cluster numbers are usually not accurate, causing low performance in the following clustering. Another strategy is to select optimal cluster numbers according to the clustering results. For example, IKAP [20] clusters cells by overestimating the cluster number in the PC space of Principle Component Analysis (PCA) [21], and then iteratively merges the nearest clusters to determine the optimal cluster number. The recently developed MultiK [22] generates multiple groups of clustering results using different cluster numbers by tests and trials and selects the optimal cluster number under certain evaluation criteria. Similar strategies are applied in methods like Clustree [23], scClustViz [24], and TooManyCells [25]. Nevertheless, these methods are machine learning or statistics-based methods that have to decouple the feature extraction and clustering into two separate steps, whereas the pre-extracted features are not optimal for the subsequent clustering. At the same time, they learn cell representations through linear algorithms (mainly PCA), which cannot efficiently process the complex scRNA-seq data [26]. Additionally, since these algorithms need multiple tests and trials, they are time-consuming, and can’t process large datasets with thousands of cells.

Here, we proposed ADClust, an automatic deep embedding clustering method for scRNA-seq data, which can accurately cluster cells without requiring a predefined cluster number. Specifically, we first pre-train the autoencoder to learn the non-linear low-dimensional representation of original gene expression, which is used to cluster cells into a mass of micro-clusters. The micro-clusters are then compared in between through a statistical test for unimodality called Dip-test [27] to detect similar micro-clusters, and similar micro-clusters are merged through jointly optimizing the carefully designed clustering and autoencoder loss functions. This process continues until convergence. By benchmarked on 11 real scRNA-seq datasets, ADClust was shown to outperform existing methods in terms of both clustering performance and the accuracy on the number of the determined clusters. More importantly, ADClust showed a high speed and scalability on large datasets.

## 2. Materials and Methods

### 2.1 Datasets and pre-processing

We employed the commonly used datasets from ref [28] that included 15 datasets. By removing seven small datasets containing less than 1000 cells, we finally kept eight datasets (Xin, Tasic, Baron Mouse, Klein, Romanov, Zeisel, Segerstolpe, and Baron Human). In order to test the scalability of our model, we selected three largest datasets (Mouse retina, TM, and PBMC 68K) from previous studies [11, 29], containing 27,499, 54,865, and 68,579 cells, respectively. As detailed in Table 1, these datasets are involved in different biological processes and various tissues and contain different scales of cells from thousands to tens of thousands derived from various single-cell RNA-seq techniques. Each dataset was preprocessed using the standard procedure as proposed in Seurat. Concretely, we normalized the gene expression by the “NormalizeData” function with the default parameter “LogNormalize” and the scaling factor of 10,000. Then, the top 2000 highly variable genes were selected through the “FindVariableFeatures” function based on the normalized matrix.

**Table 1.**
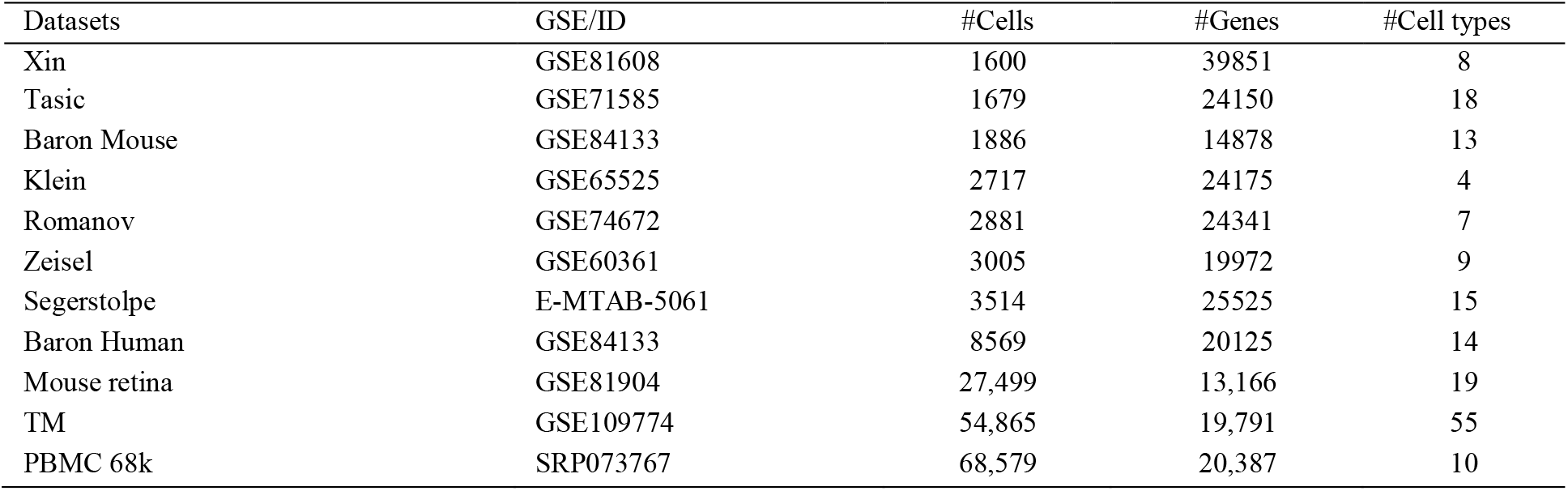
Summary of the datasets used in this study

### 2.2 The architecture of ADClust

This study proposed an automatic deep embedding clustering method that can accurately cluster cells without requiring to predefine the number of clusters. As shown in Fig. 1, the ADClust model consists of two modules: the autoencoder and clustering modules. The autoencoder aims to learn deep embedding representations of cells, and the clustering module uses the learned embedding representations to cluster cells.

**Fig 1.**
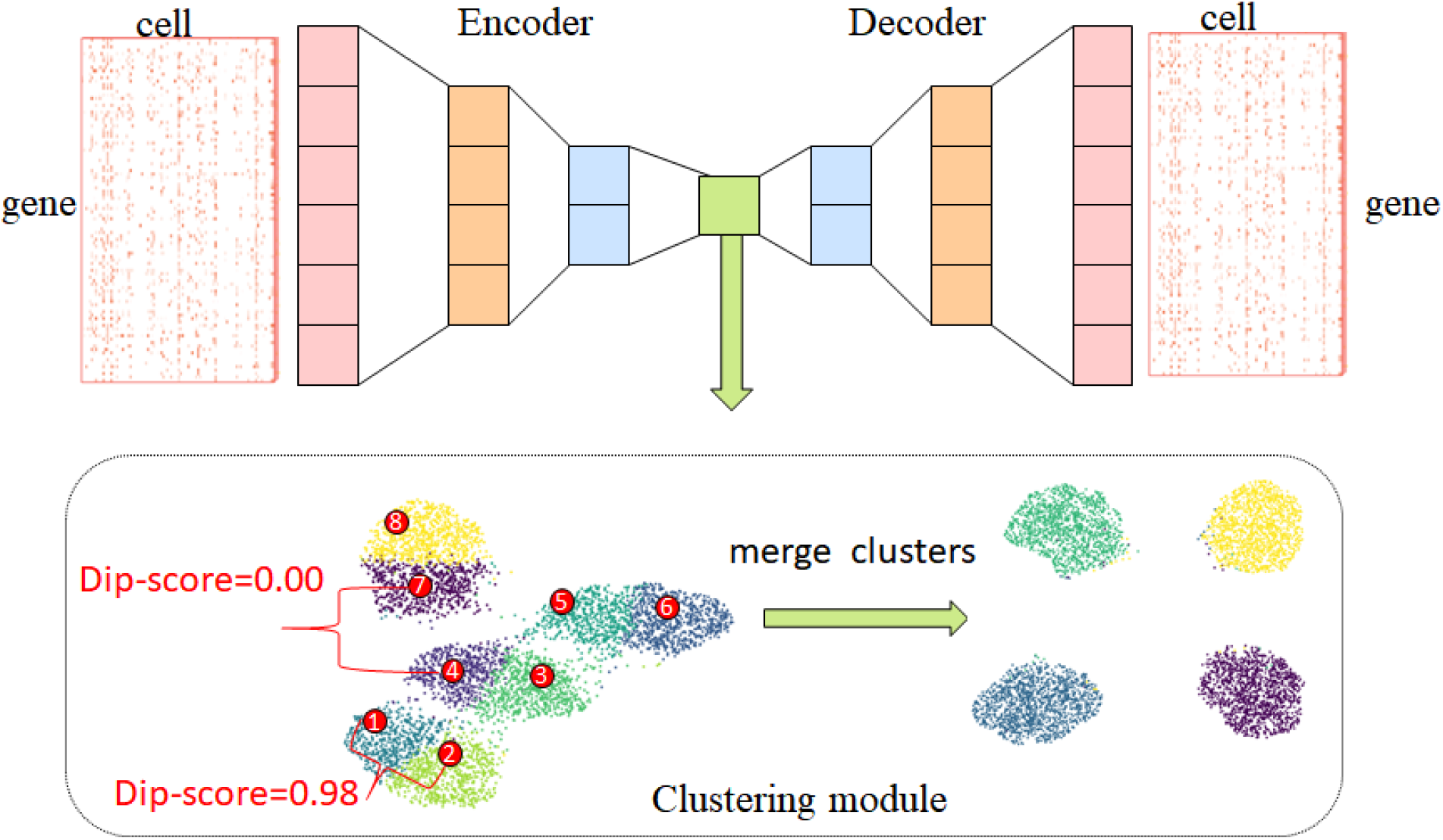
Overview of the ADClust framework. ADClust consists of autoencoder and clustering modules. The autoencoder aims to learn deep embedding representations of cells, and the clustering module uses the learned embedding representations to cluster cells.

#### 2.2.1 Autoencoder module

Autoencoder is used for embedding the input scRNA-seq gene expression data X ∈ ℝ ^*N* x *D*^ into low-dimensional space, where and are the number of cells and the size of genes, respectively. Autoencoder is an unsupervised neural network that consists of the encoder and decoder modules [30]. The encoder tries to embed the input data into a latent, and the decoder tries to reconstruct the embedded data into its origin space. Thus, the autoencoder can efficiently learn the useful low-dimensional latent by minimizing the reconstruction loss L_*res*_ as follows:

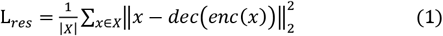

where 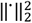 represents the square Euclidean, and the dec() and enc() represent encoder and decoder functions, respectively. The *enc*(*x*) is the learned embedding representation for gene expression *x* of individual cell.

#### 2.2.2 Clustering module

Based on the learned embedding representations, cells are clustered through the Louvain algorithm [31] into plentiful initial micro-clusters. The micro-clusters are then compared in between by Dip-test, and similar micro-clusters are merged through a carefully designed clustering loss function.

##### Clustering loss

Similar micro-clusters were pulled together in the embedding space of autoecnoder through the clustering loss L_*clu*_ originally used in ref [32]:

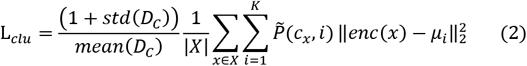

where c_*x*_ is the cluster containing cell *x, µ* is the centres for *K* clusters,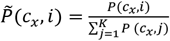 reflects cluster similarity by normalizing the Dip-score *P*(*c*_*x*_,*j*) between the cluster centres of c_*x*_ and *i* estimated through the Dip-test[27]. The “mean” and “std” are the mean and standard deviation of the set of cluster-pairwise distances *D*_*C*_.

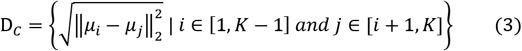

Intuitively, our model will minimize the loss by reducing the distance between a cell to the centre of its assigned centre. At the same time, the cell will also be pulled toward its similar clusters with strength depending on the similarity to the cluster *i*. As a result, the model will reduce the distance between clusters if they are similar with a large Dip-score. This process will pull similar micro-clusters together. The division by the mean (D_*C*_) was used to prevent the autoencoder from only reducing the embedding scale to minimize loss L_*clu*_. We also apply the term std (D_*C*_)to impede the model’s ability to reduce the scale and simultaneously push individual clusters far away.

Finally, we optimize ADClust with the following loss in an end-to-end manner:

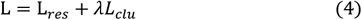

where λ is the hyper-parameter to balance contributions from the clustering loss. In this study, we set λ = 1 for all datasets.

##### Merging process

We merge two clusters if their corresponding Dip-score is larger than the Dip-score threshold. Cells in these two merging clusters will be assigned the same cell label, and a new centre of these cells will be computed as the following:

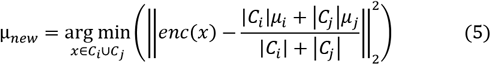

Based on the new centre µ_*new*_, we need to update the Dip-score matrix P and 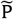. The merging process is repeated if there is still a Dip-score in *P* that is greater than the threshold. After the merging process, we continue optimizing the model and merging the clusters. This process continues until there is no Dip-score greater than the threshold.

### 2.3 Hyper-parameters setting

The ADClust was implemented in PyTorch and C. The dimensions of the autoencoder were set to input-512-256-128-10-128-256-512-input. The training batch size was generally set as 128, while the size was increased for large datasets (1024 for above 10,000 cells) to further reduce the training time of each epoch. The models were optimized through the Adam optimizer with a learning rate of 0.0001. We empirically set resolution=3 in the Louvain algorithm for all datasets to obtain the initial cluster numbers that were much larger than the true cluster numbers (We listed the true cluster numbers and the initial cluster numbers for all datasets in Supplementary Table S1). The Dip-score threshold was set to 0.9 that determined whether two clusters should be merged. The number of epochs for the pre-training and the clustering process was set to 100 and 50, respectively. All results reported in this paper were conducted on Ubuntu 16.04.7 LTS with Intel® Core (TM) i7-8700K CPU @ 3.70 GHz and 256 GB memory, with the Nvidia Tesla P100 (16G).

### 2.4 Evaluation criteria

Three common clustering metrics are used for evaluating cell clustering results in this study, Normalized Mutual Information (NMI) [33], Adjusted Rand Index (ARI) [34], and Clustering Accuracy (CA) [35]. The NMI is defined as:

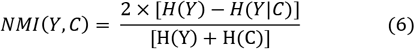

where C and Y are the predicted clusters and the true clusters (the same below), respectively. The term *H* () is used for computing the entropy.

The ARI is defined as:

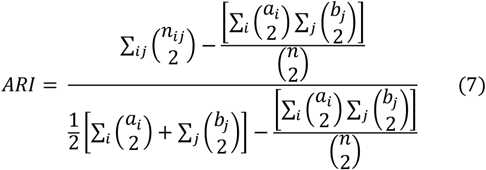

where a_*i*_ is the number of cells appearing in the i-th cluster of C, b_*j*_ is the number of cells appearing in the j-th cluster of Y. n_*ij*_ is the number of overlaps between the j-th cluster of Y and the i-th cluster of C.

The CA is calculated as:

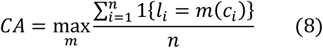

where *n* is the total number of cells, and *m* ranges over all probable one-to-one mapping between clustering assignment *c*_*i*_ and real label *l*_*i*_.

### 2.5 Benchmark methods

To evaluate the clustering performance, we compared ADClust with other tools including MultiK, SIMLR, scDeepCluster, SC3, scQcut [36], IKAP, CIDR [37], Seurat (version 3.0), and DESC. As MultiK outputs multiple estimated cluster number, we selected the estimated cluster number with the highest ARI. We set the “NUMC” parameter of SIMLR as a range [2:20] to estimate the cluster number followed ref [36]. We set the true cluster numbers for scDeepCluster since it could not estimate the cluster numbers. For other competing methods, we used the default hyper-parameters recommended in the origin paper to estimate the cluster numbers

## 3. Results

### 3.1 Performance on scRNA-seq datasets

To evaluate the clustering performance of ADClust, we applied our model to eleven scRNA-seq datasets, including eight small datasets (containing less than 10,000 cells) and three large datasets (containing more than 20,000 cells). On the eight small datasets, our model showed superior clustering performance compared to competing algorithms (Fig. 2a). On average, ADClust achieved ARI of 0.78, which was 8% higher than the one achieved by the 2^nd^ best method MultiK. The 3^rd^ ranked method scQcut is a graph partitioning algorithm achieving ARI of 0.61. This value was 10% higher than Seurat, another graph partitioning algorithm. The better performance by scQcut is likely because scQcut optimized the number of neighbors for the KNN-graph [36]. The 4^rd^ ranked method DESC achieved decent performance since it jointly optimized cell labels assignment and learned the latent representation that was fitted for clustering. CIDR and SIMLR achieved similar and low performance since their pre-estimated cluster numbers in advance were usually incorrect. scDeepCluster ranked the ninth, although it was inputted with the real cluster numbers. This is likely because its performance heavily relied on the initialized results of K-means. SC3 performed the worst since it was sensitive to parameters used in dimension reduction and tended to overestimate the cluster numbers, as also indicated in previous studies [22, 38].

**Fig 2.**
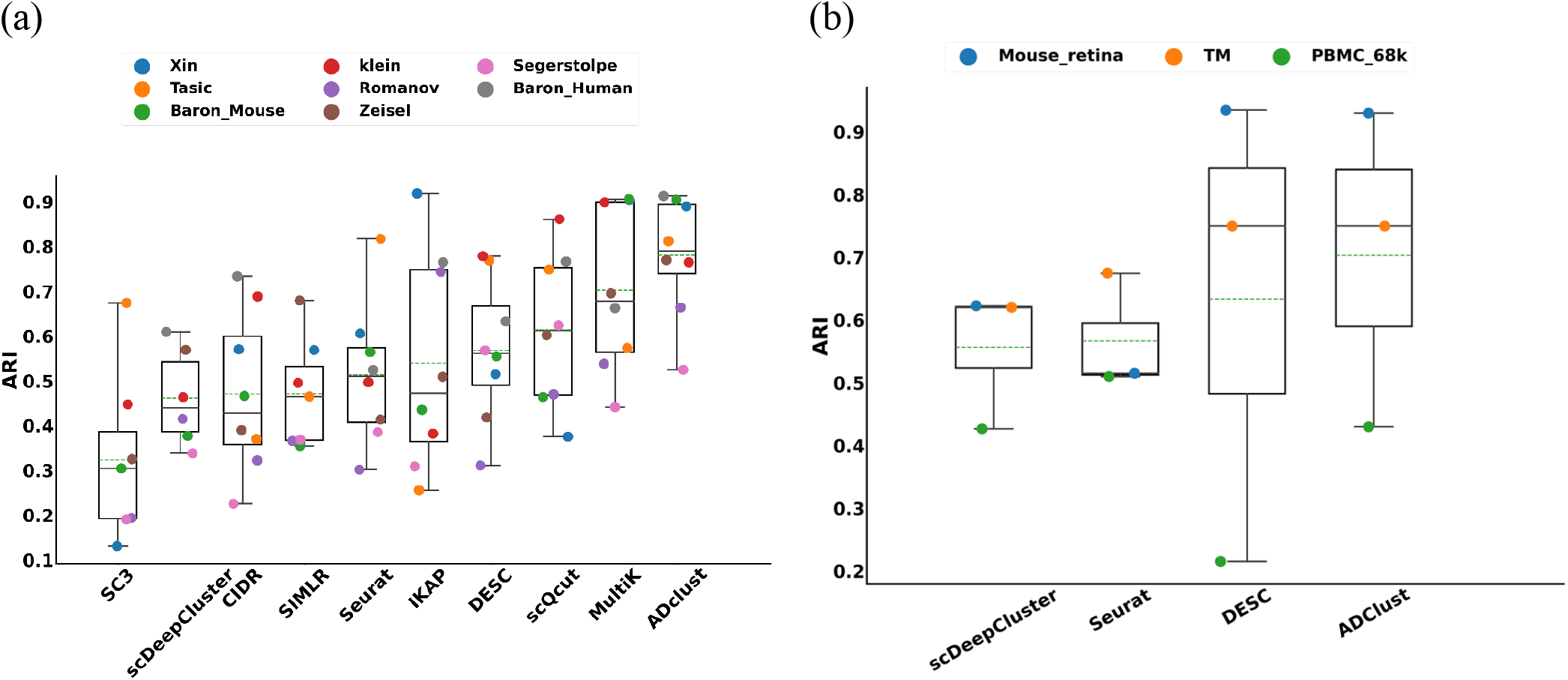
Clustering performance of all methods on (a) small scRNA-seq datasets with less than 10000 cells and (b) large scRNA-seq datasets with cells greater than 20000. The dashed line indicates the mean value of ARI.

On three large datasets with the number of cells greater than 20,000, ADClust consistently achieved the best clustering performance (Fig 2b). On average, ADClust achieved ARI of 0.70, 6% higher than the one achieved by the 2^nd^ best method DESC. The 3^nd^ ranked method Seurat achieved similar performance with the 4^nd^ ranked method scDeepCluster. IPKA and scQcut only could run on the large dataset Mouse retina, and the ARI values of them were 81% and 8% smaller than our method (ARI=0.93), respectively. We didn’t compare with IPKA and scQcut (on large datasets containing greater than 50,000 cells), CIDR, SC3, SIMLR, and MultiK due to occurring errors (scQcut), out of memory (SIMLR and CIDR), “NAN” values generation (SC3), or the runtime of more than two days (IPKA and MultiK).

We also showed the comparisons on eleven scRANA-seq datasets for evaluation criteria CA and NMI in Supplementary Figure S1, and similar trends could also be observed.

### 3.2 The evaluation on the determined cluster numbers

To evaluate the accuracy of the determined cluster number, we applied our model on all scRNA-seq datasets. Since Seurat, scDeepCluster, and DESC couldn’t estimate the cluster number, we didn’t compare with them. As shown in Fig. 3(a), on eight small datasets, the median absolute deviation of the cluster number determined by our model was closest to zero, which was the smallest of the seven methods. We further showed the specific cluster number determined by each method in Supplementary Table S2. For the eight small datasets, the cluster numbers determined by our model in five datasets were the most accurate. scQcut achieved second best performance and made the most accurate estimation for four datasets. IKAP achieved third best performance and made the most accurate estimation for three datasets. The cluster numbers determined by SIMLR, CIDR, and SC3 were usually incorrect. When tested on three large datasets, our model achieved more accurate estimation than IKAP and scQcut (Supplementary Table S2). Other clustering methods couldn’t achieve corresponding results on larger datasets due to error generation or the runtime of more than two days.

**Fig 3.**
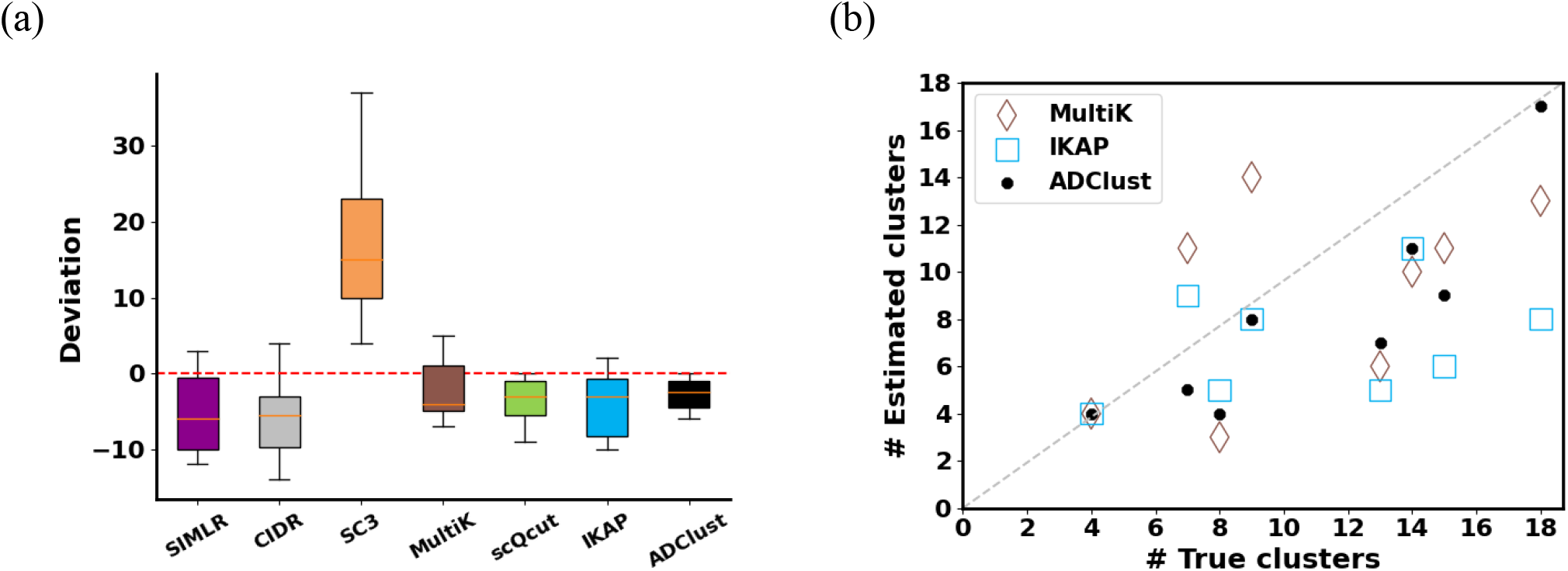
The accuracy of the determined cluster numbers. (a) the deviations between the true cluster numbers and the determined cluster numbers by each method on eight small scRNA-seq datasets. The median absolute deviation of the ADClust was the smallest of the seven methods. (b) Correlations between the determined cluster numbers by each method and the true cluster numbers.

To view the accuracy of the cluster number determined by each method more clearly, we further drew a scatter plot with the determined cluster number and the true cluster number. On eight small datasets, as shown in Fig. 3(b) and Supplementary Figure S2, the cluster number determined by our model was more similar to the true number of clusters when compared to MultiK, the method with the second-highest clustering performance. SC3 tended to overestimate the cluster number, but SIMLR and CIDR tended to underestimate the cluster number. Our model also tended to underestimate the cluster number on a few datasets.

To investigate why our model underestimated the cluster number on datasets Xin, Baron Mouse, Segerstolpe, Mouse retina, and TM, we analyzed the cell composition of these datasets. As shown in Supplementary Table S3-7, we found that these datasets contained multiple rare cell clusters [39], and these rare cell clusters that consist of less than or equal to 1.5% of the total cell population on average. (Xin < 1.5%, Baron Mouse < 0.7%, Segerstolpe <0.5%, Mouse retina <1% and TM < 0.09%). In summary, all methods, except SC3, tended to underestimate the cluster number due to the inclusion of rare cell clusters in many datasets. However, the cluster numbers estimated by SC3 were much larger than the true cluster numbers.

### 3.3 Contribution of Components to the Clustering

To investigate the contributions of components for the clustering performance of ADClust, we conducted ablation studies on all scRNA-seq datasets. As shown in Table 2, the initial clustering results of ADClust achieved the worst performance with 0.236, 0.620, and 0.347 in terms of ARI, NMI, and CA on average, respectively. The results showed that ADClust failed to achieve the desired performance when the cluster number was overestimated. We noticed the value of NMI was much greater than both ARI and CA. This is likely because initially the number of micro-clusters were much larger than the actual cluster number and each initial micro-cluster might contain only one cell type, resulting in a wrongly high NMI value. The removal of both clustering and autoencoder losses caused decreases of 7%, 6%, and 9.6% in terms of ARI, NMI, and CA, respectively. The changes indicated ADClust could achieve decant performance by jointly optimizing cell labels assignment and learning embedded representations. The removal of the clustering loss caused decreases of 6%, 3.9%, and 7.5% in terms of ARI, NMI, and CA on average, respectively.

**Table 2.**
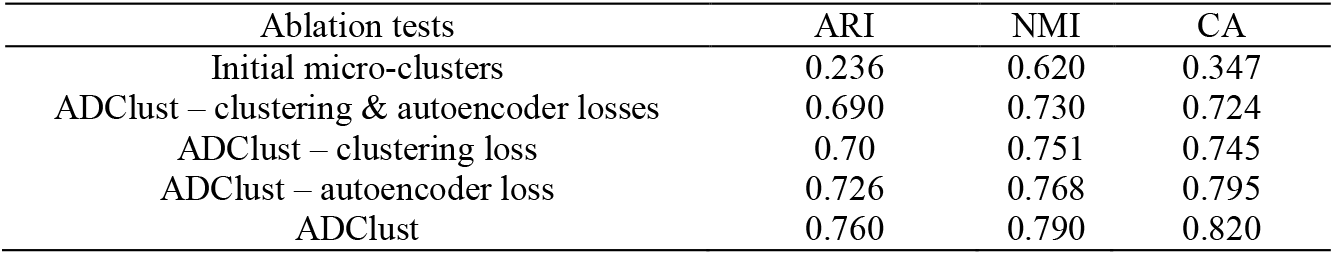
Ablation results on all datasets

The results indicated the similar micro-clusters were efficiently pulled together in the low-embedding representation of the autoencoder by optimizing clustering loss. The removal of autoencoder loss in the clustering phase caused a small but significant drop (3.4%, 2.2%, and 2.5% in terms of ARI, NMI, and CA, respectively), indicating the importance of autoencoder for improving the representation. In summary, the better clustering of the scRNA-seq data relied on the cooperation of the modules.

### 3.4 Illustration of the ADClust

To illustrate how our model worked, we visualized the merging process through UMAP [40]. Here, we took the Baron Human dataset containing 14 original cell types as an example. As shown in Fig. 4 (a), the Baron Human dataset was clustered into initial classes in this example by using the Louvain algorithm with resolution=3.0. By minimizing clustering and autoencoder loss functions, similar micro-clusters were pulled together. As shown in Fig. 4(b), most of the initial clusters were mixed with their similar clusters, resulting in multiple larger clusters with the characteristics of intra-cluster compactness and inter-cluster separability. Compared with the true cell clusters as shown in Fig. 4(c), most similar micro-clusters were correctly combined by our model. The results indicated our model could efficiently cluster cells without requiring a predefined cluster number.

**Fig 4.**
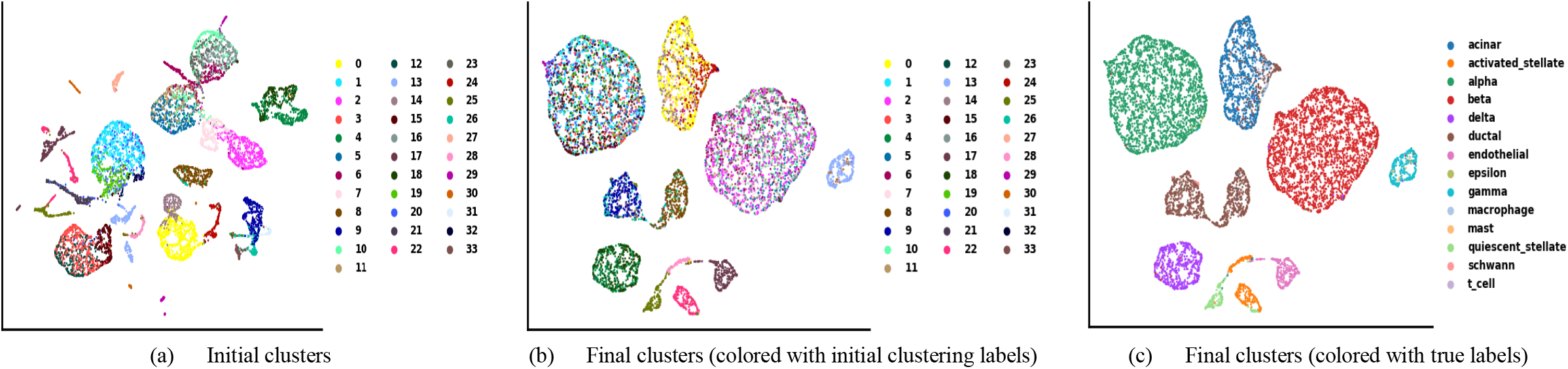
Visualizing our method on the Baron Human dataset containing 14 cell types. (a) the Baron Human dataset was clustered into initial classes by using the Louvain algorithm with parameter resolution=3.0. (b) final clustering results colored with initial clustering labels (c) final clustering results colored with true labels.

To further confirm the clustering performance of ADClust, we visualized the wrongly clustered cells by the Sankey river plots on the Baron Human dataset. As shown in Fig. 5, ADClust achieved CA, NMI, and ARI values of 0.89, 0.88, 0.913, respectively. For the two major cell types beta and alpha, which together account for the biggest portion (57%), our model could correctly assign 98% cells. The second-best method CIDR could correctly assign 93% of cells. Other methods made the accuracy of 60-82% on beta and alpha cell types (Supplementary Figure S3). One major source of wrong assignments in our model was the separation of the ductal cells into three clusters. The separation of ductal cells was also seen in all competing methods. These similar mistakes may come from the difficulty of clustering this cell type.

**Fig. 5.**
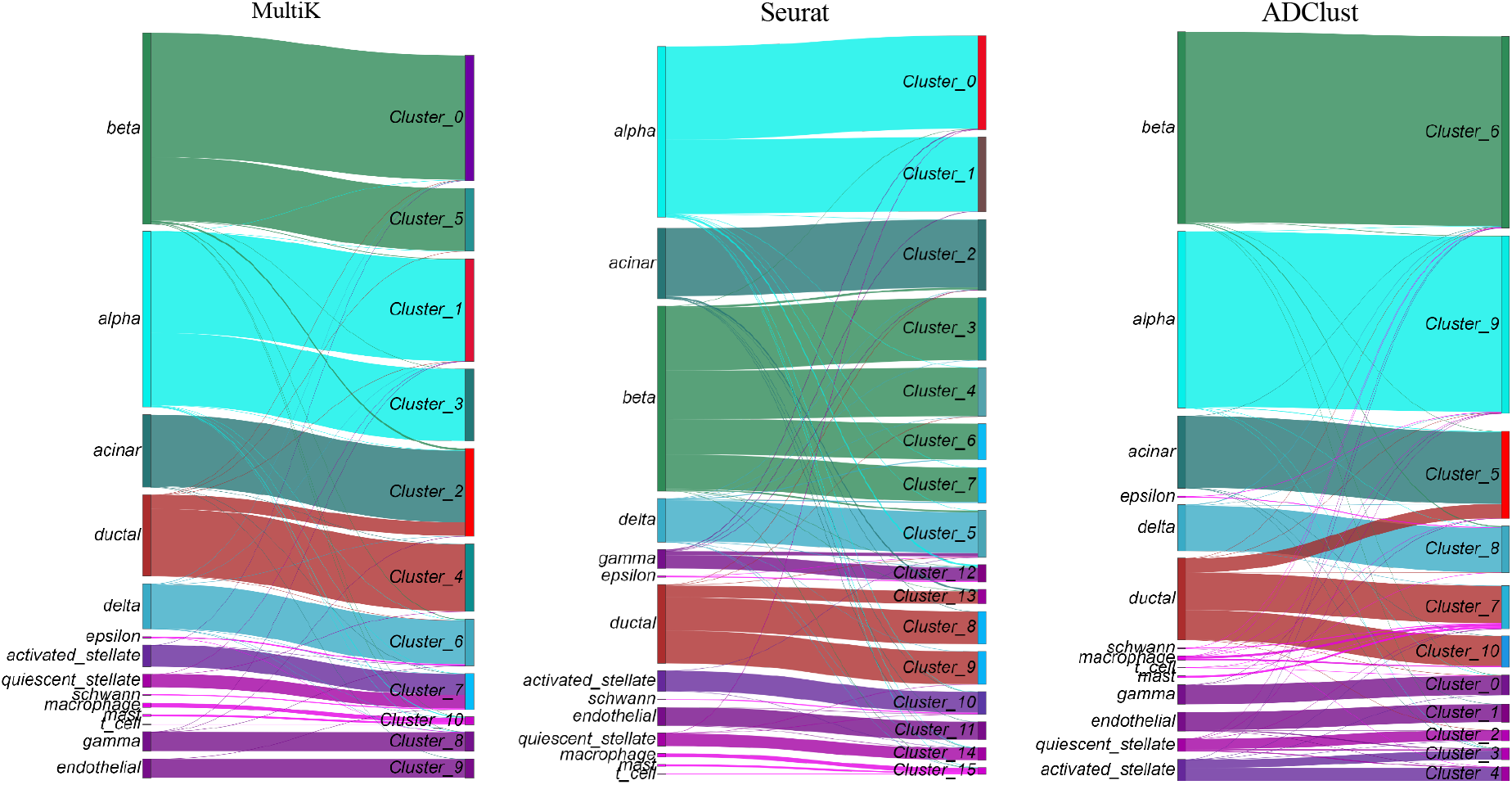
A Sankey river plot shows the match between the actual labels and clustering results on the Baron human dataset.

### 3.5 Running time

With advances in scRNA-seq technologies, the cells in emerging scRNA-seq datasets can exceed hundreds of thousands, requiring their scalability and efficiency of methods. For evaluating the runtimes of all methods and their scalability, we applied all methods to scRNA-seq datasets with a wide range of sizes. As shown in Fig. 6, dramatic differences in runtimes can be observed among these methods with increasing the number of cells. ADClust was faster than all competing clustering methods. ADClust showed high scalability with about linear growth of runtimes with the number of cells: 36s for about 2K cells and 900s for about 70K. The next fastest method CIDR was close to our algorithm in speed for datasets with less than 4K cells, but the runtimes remarkably increased with the increase in the number of cells. When the number of cells reached 8K, CIDR was > 5 times slower than our model. MultiK was the slowest method and significantly slower than all methods, which needed more than two days when running datasets with larger than 10K cells. We didn’t include partial algorithms for large datasets because they failed to run due to out of memory (SIMLR and CIDR) or “NAN” values generation (SC3) or the runtime of more than two days (IKAP and MultiK). Though Seurat was faster than our model, its ARI was averagely 23% lower than our method (Supplementary Figure S4).

**Fig. 6.**
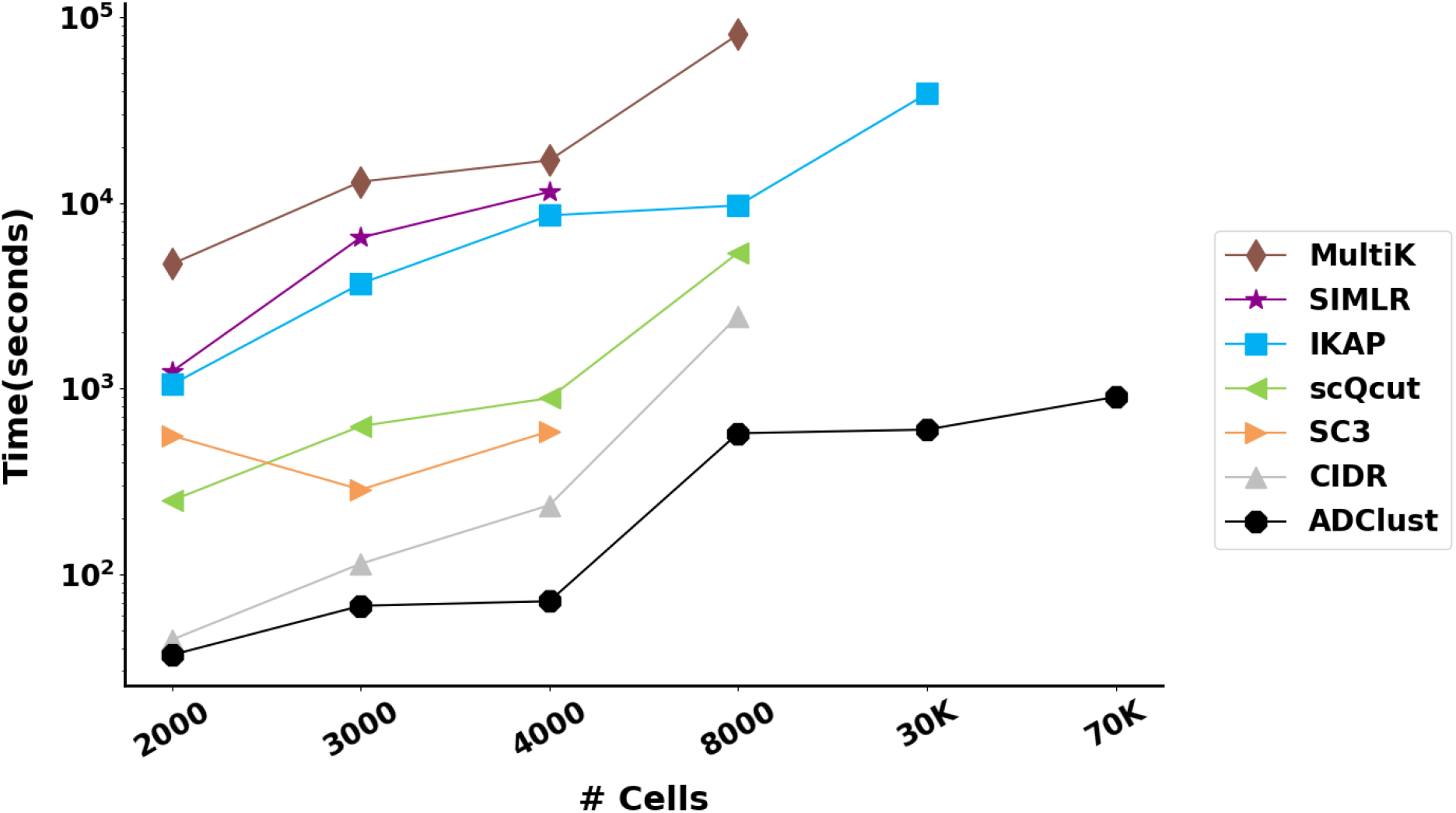
Comparison of different methods for the running time on variably sized datasets.

## 4. Discussion

The optimization of clustering algorithms is being consumingly studied in scRNA-seq analysis. One critical challenge of clustering algorithms is to accurately cluster cells into meaningful groups without predefining the cluster number. For this challenge, we proposed ADClust, an automatic deep embedding clustering method for scRNA-seq data, which can accurately cluster cells without requiring a predefined cluster number. ADClust first clusters cells into the overestimated number of micro-clusters and then pushes micro-clusters sharing structural similarities together by jointly optimizing the clustering and autoencoder loss functions. On 11 real scRNA-seq datasets, our model demonstrated better performance in terms of both clustering performance and the accuracy on the number of the determined clusters. More importantly, our model provided high speed and scalability for large scRNA-seq datasets.

While a few methods, such as MultiK, are also used for simultaneously clustering scRNA-seq data and estimating the cluster number through multiple tests and trials, it’s necessary to strike a balance between performance and time consumption for these methods. In contrast, we cluster cells by iteratively merging similar micro-clusters through minimizing clustering and autoencoder loss functions. Our model achieved superior clustering performance by jointly optimizing the cell labels assignment and learning the representations that are suitable for the clustering. More importantly, ADClust is scalable and fast since we train our model with the means of mini-batches by using GPU. In short, our model achieved superior results in terms of both performance and efficiency.

Despite the advantages of ADClust, our model can be improved in several aspects. First, our model may fail to distinguish between subtypes of cells since they have extremely similar gene expressions. We could add prior information such as marker genes into our model. Second, our model doesn’t consider batch effects and we will add modules to remove batch effects[14]. This is important with the decreasing scRNA-seq costs and increasing international collaborations. Third, small and rare clusters may not be detected by our model since the Dip-test might identify two clusters as unimodal if they differ greatly in sizes.

In summary, we demonstrate that ADClust provides an automatic deep embedded clustering algorithm, which provides stable clustering solutions for scRNA-seq datasets without requiring the predefined cluster number. In addition, it is worth noting that the concept of ADClust is applicable beyond scRNA-seq data, such as mass cytometry and scATAC-seq data.

## Code availability

All source codes used in our experiments have been deposited at https://github.com/biomed-AI/ADClust.

## Data availability

The scRNA-seq datasets that support the findings of this study are available here: https://www.synapse.org/#!Synapse:syn26524750/files/.

